# Neonicotinoids Disrupt Circadian Rhythms and Sleep in Honey Bees

**DOI:** 10.1101/2020.04.15.042960

**Authors:** Michael C. Tackenberg, Manuel A. Giannoni-Guzmán, Caleb A. Doll, José L. Agosto-Rivera, Kendal Broadie, Darrell Moore, Douglas G. McMahon

**Affiliations:** Vanderbilt Brain Institute, Vanderbilt University, Nashville, TN, USA; Department of Biological Sciences, Vanderbilt University, Nashville, TN, USA; Department of Biology, University of Puerto Rico – Río Piedras, San Juan, PR, USA; Department of Biological Sciences, East Tennessee State University, Johnson City, TN, USA

## Abstract

Honey bees are critical pollinators in ecosystems and agriculture, but their numbers have significantly declined. Declines in pollinator populations are thought to be due to multiple factors including habitat loss, climate change, increased vulnerability to disease and parasites, and pesticide use. Neonicotinoid pesticides are agonists of insect nicotinic cholinergic receptors, and sub-lethal exposures are linked to reduced honey bee hive survival. Honey bees are highly dependent on circadian clocks to regulate critical behaviors, such as foraging orientation and navigation, time-memory for food sources, sleep, and learning/ memory processes. Because circadian clock neurons in insects receive light input through cholinergic signaling we tested for effects of neonicotinoids on honey bee circadian rhythms and sleep. Neonicotinoid ingestion by feeding over several days results in neonicotinoid accumulation in the bee brain, disrupts circadian rhythmicity in many individual bees, shifts the timing of behavioral circadian rhythms in bees that remain rhythmic, and impairs sleep. Neonicotinoids and light input act synergistically to disrupt bee circadian behavior, and neonicotinoids directly stimulate wake-promoting clock neurons in the fruit fly brain. Neonicotinoids disrupt honey bee circadian rhythms and sleep, likely by aberrant stimulation of clock neurons, to potentially impair honey bee navigation, time-memory, and social communication.

## Introduction

Honey bees are critical pollinators in ecosystems and agriculture, but their numbers have significantly declined over the past three decades^1^. The observed declines in pollinator populations are thought to be due to multiple factors including habitat loss, climate change, increased vulnerability to disease and parasites, and pesticide use^2^. Neonicotinoid pesticides are agonists of insect nicotinic cholinergic receptors, which mediate excitatory neurotransmission within the insect brain, and sub-lethal exposures to neonicotinoids are linked to reduced honey bee hive survival^3,4^.

Honey bees are highly dependent on circadian clocks to regulate critical behaviors, such as foraging orientation and navigation, time-memory for food sources, sleep, and learning/ memory processes^5–8^. Circadian clock neurons in *Drosophila* receive light input through cholinergic signaling^9^, and honey bees possess homologous clusters of putative clock neurons^10,11^, thus honey bee clock neurons are potential targets for neonicotinoid effects. Sub-lethal doses of neonicotinoid pesticides have been shown to disrupt navigation by forager honey bees^12,13^. In addition, neonicotinoid pesticides disturb honey bee learning and memory^14,15^ The mechanisms by which neonicotinoids disrupt honey bee orientation and learning are unknown, however both of these complex behaviors are highly regulated by endogenous honey bee circadian clocks^8,16^. Sleep is an additional critical clock-regulated neural process, and sleep in honey bees supports navigational memory consolidation^17^ and social communication through the waggle dance by which foragers communicate the location of food sources^18^.

Here we tested for effects of ingestion of neonicotinoids through feeding on honey bee circadian rhythms and sleep. We find that neonicotinoid ingestion over several days results in neonicotinoid accumulation in the bee brain, disrupts circadian rhythmicity in many individual bees, shifts the timing of behavioral circadian rhythms in bees that remain rhythmic, and impairs sleep. Neonicotinoids, thus, potentially impair key clock-based aspects of honey bee navigation, time-memory, and social communication.

## Results

Honey bee (*Apis mellifera ligustica*) hives were maintained on the Vanderbilt University campus. Vanderbilt does not use neonicotinoid pesticides, and naïve forager bees from our hives did not show detectable levels of thiamethoxam or clothianidin, the two neonicotinoids used in this study (Supplemental Figure 1).

To study effects of neonicotinoid pesticides on individual honey bee circadian rhythms and sleep, forager honey bees were captured at hive entrances and maintained in individual infrared activity monitors in the laboratory for up to 8 days with *ad libitum* access to bee candy (honey and powdered sugar, with or without the addition of thiamethoxam or clothianidin)^19^. Neonicotinoids in the food ranged from 0-140 ng/g (or parts per billion, ppb), which is within the concentration range reported in flower nectar and pollen encountered by foraging bees^20–23^. To determine the amounts of neonicotinoids present in dosed bees, we isolated whole brains from control bees that consumed bee candy with no added neonicotinoid, and from bees that consumed bee candy dosed with thiamethoxam at two concentrations (25 ppb and 70 ppb). We collected brains from each group at 24-hour intervals over 4 days and then tested for thiamethoxan and its metabolite clothianidin by mass spectrometry (see Methods). Thiamethoxam and clothianidin were undetectable in control bee brains (Supplemental Figure 1). Thiamethoxam was undetectable, or detected at low picogram levels, in bee brains at both doses. The active metabolite of thiamethoxam, clothianidin^24,25^, was readily detected at higher levels (0-220 pg/brain, Supplemental Figure 1), consistent with the previously described rapid conversion of thiamethoxam to clothianidin in insects^24,25^. Forager honey bees exposed to thiamethoxam-dosed food exhibited sustained levels of clothianidon of ca. 50-70pg/brain through day 4 at the 70 ppb dose, with lower levels detected at the 25 ppb dose (Supplemental Figure 1).

To obtain sufficient circadian cycles for analysis, only data from control and dosed bees that survived a minimum of 4 days in light:dark (LD) experiments, or a minimum of 5 days in constant dark (DD) and constant light (LL) experiments, were included in datasets (see Methods). For those bees meeting these criteria for inclusion, survivorship analysis indicated that there was no additional mortality of neonicotinoid-exposed bees compared to control unexposed bees at any of the neonicotinoid concentrations (25-140 ppb) used in the LD, DD, and LL datasets (N = 944, Supplemental Figure 2). Survivorship analysis of all bees that began LD, DD and LL experiments (including those that did not survive sufficient days to contribute to the dataset) showed that there was an initial significant loss of dosed bees compared to controls primarily limited to the first day of exposure to both 70 ppb and 140 ppb levels of thiamethoxam or clothianidin in the food, followed by comparable survivorship to controls on subsequent days (Supplemental Figure 3). Thus, our study reports results from bees for which consumption of food with these levels of neonicotinoids for 4-5 days was not lethal and did not reduce survivorship compared to controls, while consumption of food with the 70 ppb and 140 ppb levels of thiamethoxam and clothianidin was lethal to some bees in the first 24 hours.

We first measured entrained and free-running circadian rhythms of captured foragers for 4 days of 12:12 LD followed by 4 days of DD in the presence and absence of neonicotinoids in the food. Ingestion of neonicotinoids completely disrupted locomotor circadian rhythms in a significant portion of bees, inducing arrhythmic behavior in an increasing proportion of bees in a dose-dependent manner (Figure 1A, B). While only a small fraction (12%) of unexposed control bees exhibited arrhythmic locomotor behavior, nearly half of thiamethoxam-dosed (46%) and clothianidin-dosed (42%) bees lost circadian behavioral rhythms following consumption of these neonicotinoids (140 ppb, Figure 1E, F; Pearson Chi-Square test, *p* = 0.0043 Thiamethoxam; *p* = 0.0142 Clothianidin).

**Figure 1.**
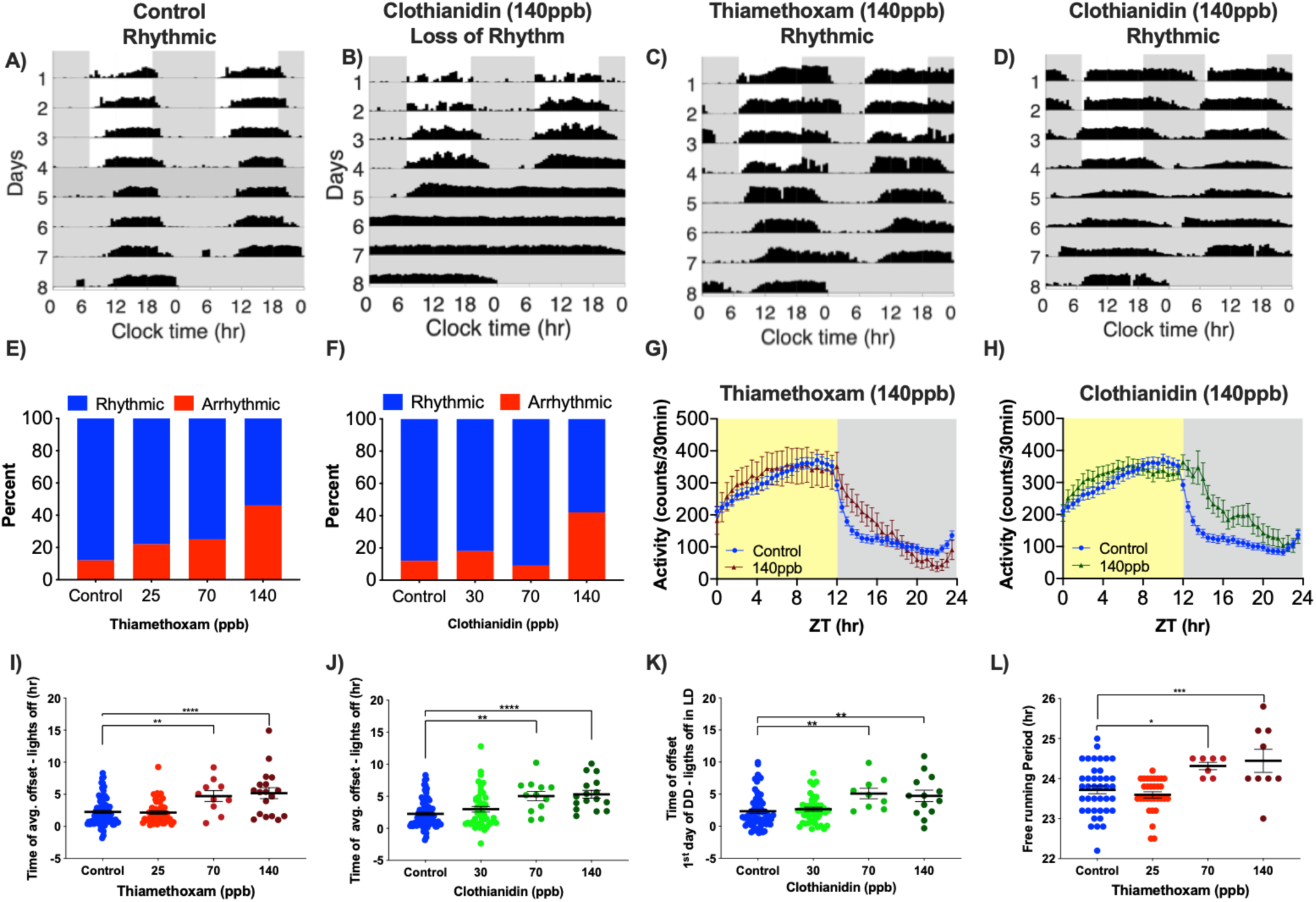
Neonicotinoid ingestion alters circadian locomotor rhythms of honey bee foragers in LD and DD. Representative actograms of forager bees showing **A)** control rhythmic activity, **B)** loss of rhythms following ingestion of clothianidin (140 ppb) and altered locomotor rhythm patterns after ingestion of **C)** thiamethoxam (140 ppb) and **D)** clothianidin (140 ppb). Contingency plots reporting the percent of arrhythmic individuals for bees exposed to **E)** thiamethoxam (Pearson X^2^ = 13.14; *p* = 0.0043, **) or **F)** clothianidin (Pearson X^2^ = 10.59; *p* = 0.0142, *). 30min binned activity profiles of control bees vs. bees exposed to either 140 ppb of **G)** thiamethoxam (Two-way RM ANOVA, Time *p* < 0.0001, ****; Dose *p* = 0.7150; Interaction *p* < 0.0001, ****) or **H)** clothianidin (Two-way RM ANOVA, Time *p* < 0.0001, ****; Dose *p* = 0.1014; Interaction *p* < 0.0001, ****). Average time of activity offset following lights off for **I)** thiamethoxam-exposed bees (One-way ANOVA F = 11.23; *p* < 0.0001, ****; Dunnett’s multiple comparison 70 ppb *p* = 0.0054, **; 140 ppb *p* < 0.0001, ****) and **J)** clothianidin-exposed bees (One-way ANOVA F = 9.664; *p* < 0.0001, ****; Dunnett’s multiple comparison 70 ppb *p* = 0.0012, **; 140 ppb *p* < 0.0001, ****). **K)** Offset of activity during the first day of constant darkness relative to time of lights off in previous LD for clothianidin-exposed bees (One-way ANOVA F = 5.805; *p* < 0.0009, ***; Dunnett’s multiple comparison 70 ppb *p* = 0.0077, **; 140 ppb *p* = 0.0063, **). **L)** Free-running period of bees under constant darkness after 4 days of LD and ingestion of different doses of thiamethoxam (One-way ANOVA F = 7.323; *p* < 0.0002, ***; Dunnett’s multiple comparison 70 ppb *p* = 0.0374, *; 140 ppb *p* = 0.0028, **). Dunnett’s multiple comparisons were performed, only significant values are shown for panels **I-L**.

In addition to disrupting honey bee locomotor circadian rhythms in many individuals, neonicotinoid ingestion also dramatically altered the rhythmic behavior of the dosed bees that remained rhythmic. Individual and average actogram activity profiles show that honey bees that ingested neonicotinoids delayed termination of their active period well into the dark phase following lights off, and increased their activity at night (Figure 1C, D, G, H). There was a more than 3-hour delay in the time of activity offset in bees that consumed either thiamethoxam or clothianidin compared to undosed bees (Figure 1I, J; One-Way ANOVA *p* < 0.0001). This marked delay in activity offset persisted into the first day of DD in bees that consumed clothianidin (Figure 1K, *p* = 0.0007), indicating a shift in the alignment of circadian rhythms relative to the light cycle. While honey bees that consumed neonicotinoids had statistically significant increases in the proportion of locomotor activity at night, and in the overall duration of activity each day, there was no significant increase in the total activity counts, indicating that neonicotinoids induced changes in the overall temporal structure of activity, but not the total amount of daily activity (Supplemental Figure 4). In addition, ingestion of thiamethoxam, but not clothianidin, lengthened the free-running period of honey bee locomotor rhythms measured in DD following LD (Figure 1L; One-Way ANOVA, *p* = 0.0002), demonstrating an effect on a core property of the underlying circadian clock.

Surprisingly, when honey bees were exposed to neonicotinoids solely in DD (without first being exposed during LD), the treatment effects were blunted. There was no significant increase in behavioral arrhythmicity (Figure 2A,C) and only a trend toward lengthening of the period (Supplemental Figure 5). This result, and the central role of cholinergic signaling in clock light input in *Drosophila*, suggested to us the possibility of a key interaction between light input and neonicotinoid ingestion. To examine this potential interaction further, we combined continuous background illumination (LL, 250 lux) with exposure to thiamethoxam-dosed bee candy for a period of 5 days (Figure 2B,D). Constant illumination is an abnormal signal to the circadian clock which renders some bees arrhythmic^26^. We found that that LL itself disrupted rhythms in 28% of undosed bees compared to just 9% arrhythmicity in DD. When thiamethoxam was combined with LL, a dose-dependent pattern in circadian disruption emerged, such that 34% of bees exposed to 25 ppb and 46% of bees exposed to 70 ppb became arrhythmic (Fig. 2B,D), but no significant effects occurred in response to exposure to these doses of thiamethoxam in DD (Figure 2A). The differential effects between neonicotinoid exposure in LL vs DD suggests synergy between continuous light input and neonicotinoids in altering honey bee circadian behaviors.

**Figure 2.**
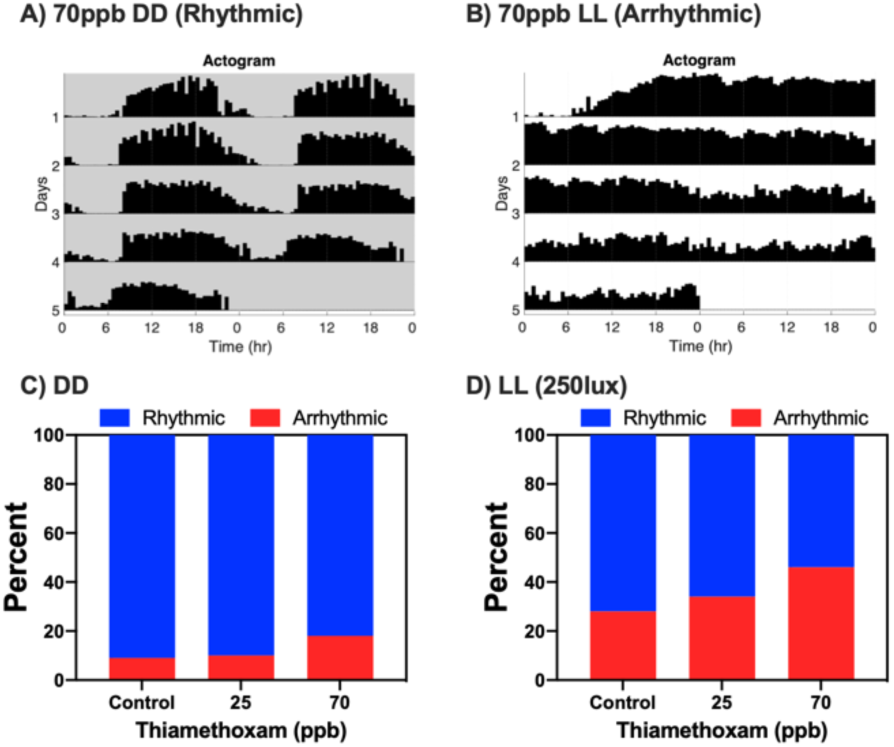
Constant light and thiamethoxam ingestion act synergistically to disrupt circadian rhythmicity of honey bees. **A)** Representative actogram of foragers showing rhythmic activity of a bee exposed to 70ppb thiamethoxam in constant darkness (DD). **B)** Representative actogram of arrhythmic behavior after exposure to 70ppb thiamethoxam in constant light (LL). Contingency plots showing the percentage of honey bees that were arrhythmic after exposure to thiamethoxam at various concentrations **C)** under constant darkness (DD) (X^2^ = 2.101 *p* = 0.3498, ns), or **D)** constant light (LL) (250 lux; X^2^ = 12.16 *p* = 0.0023, **).

Light input mediated by nicotinic cholinergic neurotransmission targets the large ventral lateral neurons (l-LN_v_s) a specific subset of PDF-expressing clock neurons that antagonize sleep in fruit flies^27,28^. We therefore tested for neonicotinoid effects on bee sleep using previously established methods for sleep analysis in flies and bees that define sleep as episodes of inactivity lasting 5 minutes or more^29–31^. Ingestion of neonicotinoids significantly impaired sleep in honey bees (Figure 3), with average daily sleep profiles showing that neonicotinoid-dosed bees in LD exhibited reduced sleep across both night and day (Figure 3A, B). Overall sleep duration was reduced by up to 50% (Figure 3C, D, One-way ANOVA *p* = 0.0004 Thiamethoxam; *p =* 0.0067 Clothianidin), while the number of sleep bouts were reduced as much as 24% (Figure 3E, F, One-way ANOVA, *p* = 0.0314 Thiamethoxam; *p* = 0.0549 Clothianidin). Neonicotinoids also significantly reduced sleep upon application in DD and in LL (Supplemental Figure 6). Thus, while neonicotinoids did not change the overall levels of activity (above), they changed the temporal structure of both activity and rest, disrupting sleep.

**Figure 3.**
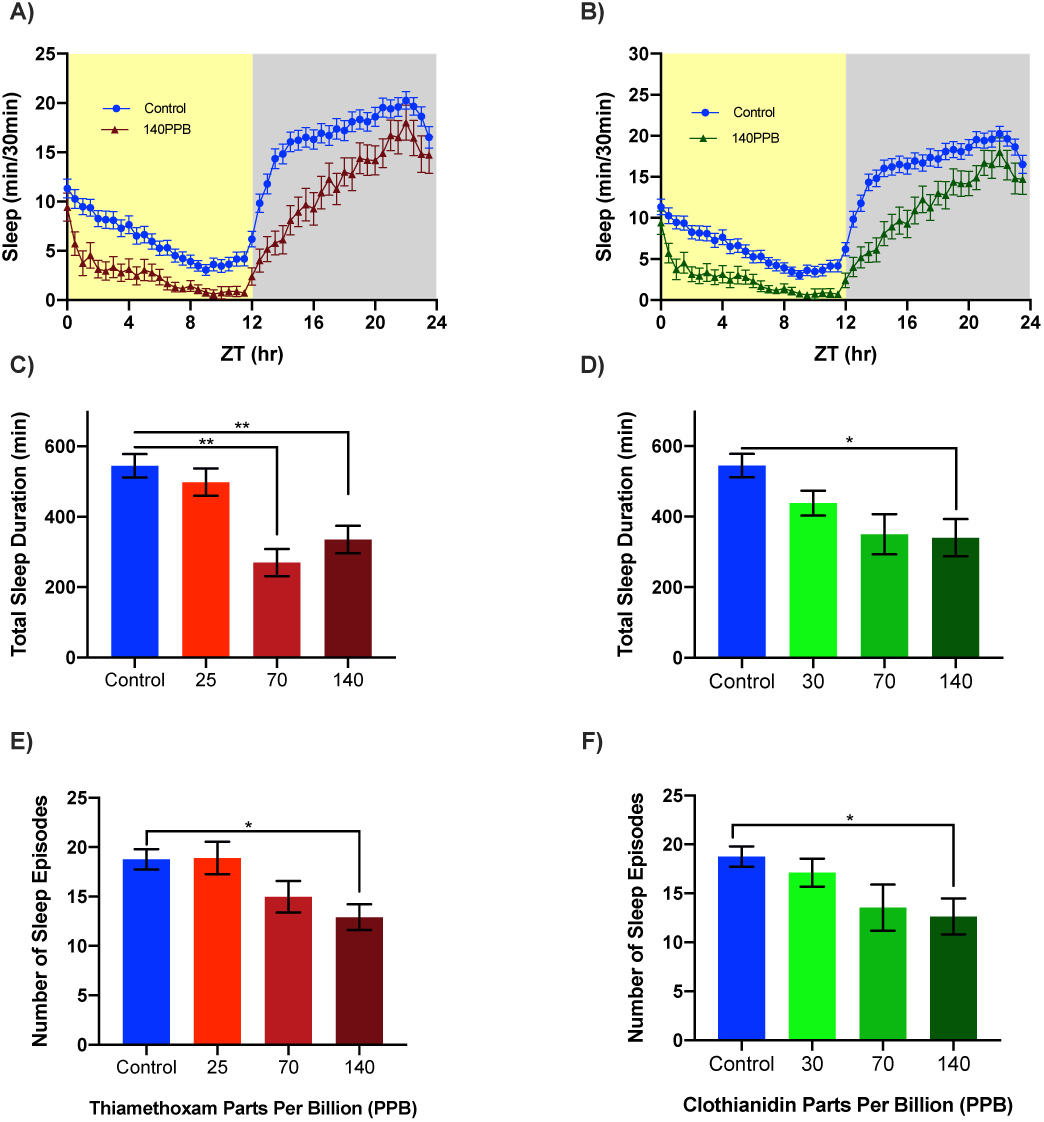
Neonicotinoids disrupt sleep in honey bee foragers. Four day average sleep profiles for control bees and bees exposed to either **A)** thiamethoxam (Two-way RM ANOVA, Time *p* < 0.0001, ****; Dose *p* = 0.0004, ***; Interaction *p* = 0.0015, **) or **B)** clothianidin (Two-way RM ANOVA, Time *p* < 0.0001, ****; Dose *p* = 0.0054, **; Interaction *p* < 0.0001, ****). Significant decreases in total sleep duration for exposure to **C)** thiamethoxam (One-way ANOVA F = 6.436; *p* = 0.0004, ***; Dunnett’s multiple comparison 70 ppb *p* = 0.0038, **; 140 ppb *p* = 0.0026, **) or exposure to **D)** clothianidin (One-way ANOVA F = 4.205; *p* = 0.0067, **; Dunnett’s multiple comparison 140 ppb *p* = 0.0115, *). In addition, there were significant decreases in the number of sleep episodes for exposure to **E)** thiamethoxam (One-way ANOVA F = 3.020; *p* = 0.0314*; Dunnett’s multiple comparison 140 ppb *p* < 0.0233, *) and for **F)** clothianidin (One-way ANOVA F = 2.586; *p* = 0.0549, ns; Dunnett’s multiple comparison 140 ppb *p* < 0.0396, *) dosed bees. Significant values only are shown for Dunnett’s post hoc multiple comparisons **C-F**.

To verify that neonicotinoids affect PDF-expressing circadian clock neurons, we next measured the Ca^2+^ responses of *Drosophila* l-LN_v_ neurons. Intact whole-brain explants expressing the GCaMP5G Ca^2+^ indicator in the PDF^+^ clock neurons were imaged for Ca^2+^-dependent changes in fluorescence intensity in the l-LN_v_ wake-promoting neurons upon exposure to either clothianidin or vehicle. We applied clothianidin because mass spectroscopy indicated an accumulation of clothianidin, as a metabolite of thiamethoxam, in bee brains when bees consumed thiamethoxam-dosed food (Supplemental Figure 1). Exposure of isolated brains to clothianidin resulted in a ca. 350% increase in Ca^2+^ fluorescence in the l-LN_v_ PDF^+^ clock neurons (ΔF/F, vehicle, −0.01 ± 0.04, Clothianidin, 3.62 ± 0.47, *p* < 0.0001, Fig. 4A, B, C). These results are consistent with findings that *Drosophila* l-LN_v_ neurons are excited directly by nicotinic cholinergic stimulation^32,33^, and likely express nicotinic acetylcholine receptors that are targeted by neonicotinoids.

**Figure 4.**
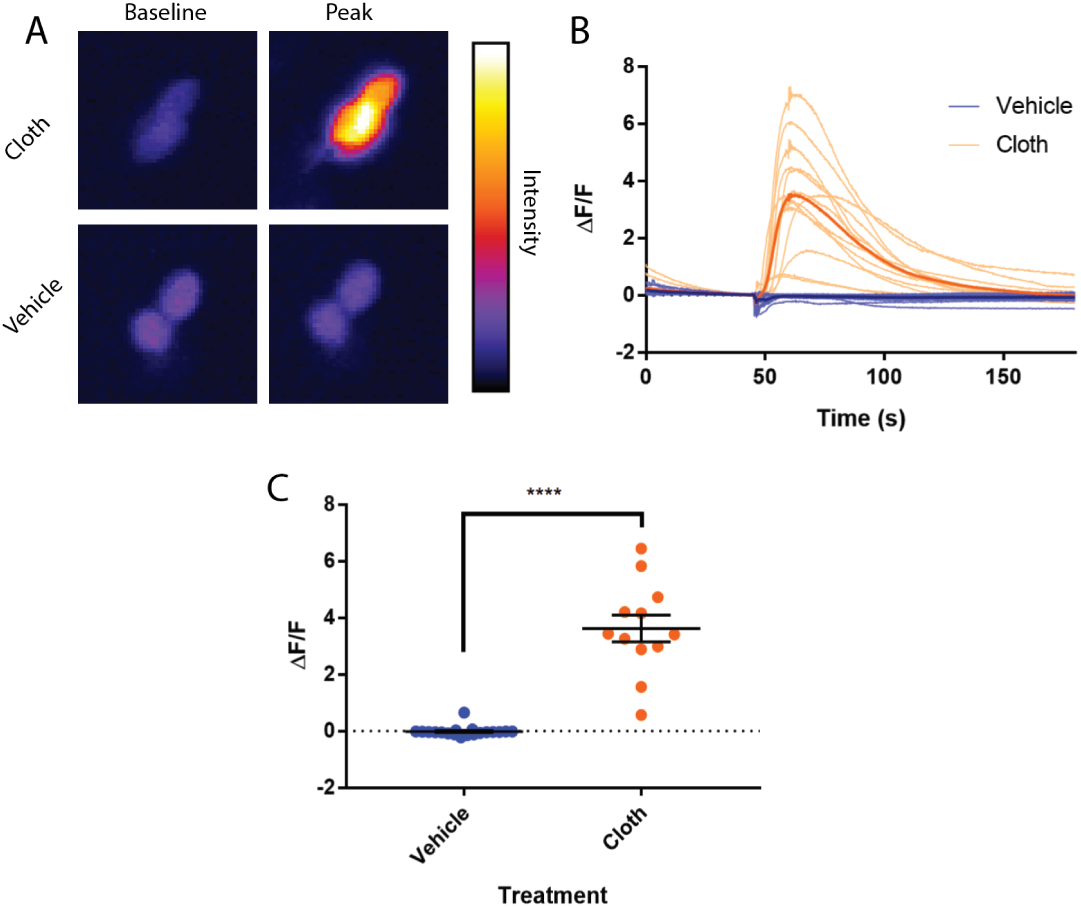
Clothianidin induces excitatory increases in [Ca^2+^]_i_ in *Drosophila* l-LN_v_ clock neurons. **A)** representative images showing the baseline and peak fluorescence in *Drosophila* PDF^+^ l-LN_v_ neurons expressing GCaMP5G fluorescent calcium indicator, in response to 4.1 ppb clothianidin (cloth, top) or vehicle (bottom). **B**) fluorescence intensity traces showing individual responses of *Drosophila* PDF^+^ l-LN_v_ neurons to clothianidin (pale orange) or vehicle (pale blue). Averaged traces are shown for explants exposed to clothianidin (orange) and vehicle (blue). **C)** average peak fluorescence responses of neurons in explants exposed to vehicle (blue) or clothianidin (orange). Error bars represent SEM. **** indicates *p* < 0.0001.

## Discussion

This study demonstrates that neonicotinoid ingestion disrupts circadian rhythms in exposed honey bees. In addition, neonicotinoids alter the rhythmic behavior of bees that remain rhythmic following ingestion, delaying activity offset and extending activity into the night. Neonicotinoids also impair honey bee sleep, reducing total sleep, and the number of sleep bouts.

The effects of neonicotinoids on honey bee circadian rhythms are enhanced by light input. In the presence of light cycles (LD) or constant light (LL), neonicotinoids induce loss of circadian behavioral rhythms in a dose-dependent manner. However, in the absence of light cycles (DD), neonicotinoids did not significantly disrupt honey bee circadian rhythms at similar exposure levels. In the field, forager honey bees experience the daily light cycle, and thus the effects of neonicotinoids measured in LD cycles are most relevant outside the laboratory.

In *Drosophila*, the PDF+ l-LN_v_ wake-promoting arousal neurons receive nicotinic cholinergic input from the compound eyes^28,34,35^. Neonicotinoid ingestion in bees mimics the effects of experimental overstimulation of *Drosophila* l-LN_v_s, extending activity into the night, increasing night-time activity, and dramatically reducing sleep^28^. Our results suggest that neonicotinoids aberrantly excite nicotinic cholinergic light input pathways to overstimulate homologous wake-promoting clock neurons in bees, leading to disruption and alteration of circadian rhythms and sleep. The neonicotinoid stimulation of *Drosophila* l-LN_v_ neurons we observed (Figure 4), and the previously observed delay of honey bee circadian rhythms in response to PDF injection into the putative bee clock nuclei^10^, support this proposed mechanism. It is also notable that chronic neonicotinoid ingestion also affects honey bee locomotor responses to light (phototaxis), and acute exposure can induce increased locomotor activity^36^.

Chronic ingestion of neonicotinoids resulted in three principal effects in our studies – disruption of behavioral circadian rhythms, shifts in the alignment of circadian activity with the light cycle, and impairment of sleep. Each of these effects on the honey bee circadian system have the potential for deleterious impacts on critical honey bee foraging behaviors which could ultimately impact hive health and survival. The abrogation of circadian rhythms that we found in significant proportions of bees that ingested neonicotinoids, could be expected to disrupt forager time-compensated sun-compass navigation, as it does in butterflies where the critical clocks are in the antennae^37^. Compromised circadian rhythms would also be expected to disrupt circadian clock-dependent time-memory for food source availability (Zeitgedächtnis). Each of these effects would significantly decrease the efficiency of honey bee foraging.

Ingestion of neonicotinoids delayed activity offset in LD, and this delay persisted when bees were then transitioned into DD, suggesting that neonicotinoids alter the timing of the honey circadian clock. The timing of activity offset relative to lights off has been shown to be a key marker of the entrainment, of honey bee circadian rhythms with the daily light cycle^38^, which is a dominant cue defining local environmental time. Thus, shifts in the alignment of circadian locomotor rhythm relative to the daily light cycle, as we observed in honey bees that remained rhythmic following neonicotinoid ingestion, would be expected to induce navigational errors due to changes in alignment of clock-based time-compensation for sun-compass orientation^16,39^. Neonicotinoid consumption induced ca. 3-hour delays in locomotor activity alignment of dosed bees compared with the light cycle. Shifts of this magnitude are predicted to engender large navigational errors of ca. 45 degrees, based on the known sun-compass time compensation of 15 degrees/hour^40^. Indeed, acute exposure to neonicotinoids impairs navigation and homing in honey bees^12,13^, as do chronic exposures^41^, and quantitative models of hive viability predict that the observed rates of forager loss due to the neonicotinoid exposure can negatively impact hive survival^12,42^.

Interestingly, catch-move-release experiments following chronic neonicotinoid exposures showed that many of the neonicotinoid-exposed bees became lost and did not return to their hive due to mis-orientation of initial vector return flight paths^41^. This deficiency is consistent with dysregulation of circadian clock dependent sun-compass orientation predicted by the neonicotinoid effects reported here. Moreover, the disruption and modification of forager honey bee circadian rhythms may affect the overall bee colony temporal structure through disruption of social entrainment. Previous studies have reported that the rhythms of honey bee foragers serve to synchronize the rhythms of the hive-bound nurse bees^43,44^.

We found that chronic ingestion of neonicotinoids impaired honey bee sleep as well as circadian locomotor rhythms. Sleep deprivation has been widely demonstrated to disrupt neuro-cognitive functions such as learning, memory, and social communication in a wide variety of species. In honey bees, sleep deprivation specifically impairs critical navigational memory consolidation, also reducing the probability of forager bees returning to the hive^17^. In addition, sleep deprivation reduces the precision of the waggle dance by which foragers communicate the location of food sources^18^, degrading the social communication critical to foraging. Taken together, the sleep-disrupting effects of neonicotinoid ingestion may also compromise key neuro-behavioral mechanisms supporting honey bee foraging and hive survival.

Our results show dramatic impacts of neonicotinoids on circadian rhythms and sleep in honey bees. However, two limitations of our studies stem from the laboratory setting in which they were performed – 1) experiments were conducted over a relatively short study period (up to about 1 week), and 2) we measured the circadian rhythms of individuals, not entire hives. The relatively short duration of these experiments may actually underestimate the impact of long-term environmental neonicotinoids on the honey bee circadian system, as in the field exposures may be more prolonged and may include developmental exposures as well. It would be of interest in the future to measure circadian aspects of whole honey bee colony behavior in the field experiments such as studies that have demonstrated reduced hive productivity and whole hive survival due to exposure to neonicotinoids^3,4^.

Overall, we find that chronic ingestion of neonicotinoids disrupts circadian behavioral rhythms and sleep in forager honey bees, and that the predicted downstream neurobehavioral impacts of disrupted rhythms and sleep are potentially deleterious to hive health. Future studies are needed to identify and characterize the fundamental neural substrates that generate circadian and sleep behaviors in honey bees, and that synchronize bee circadian rhythms to local time through the daily light cycle. The cholinergic signaling neural circuits by which light and neonicotinoids impact honey bee locomotor rhythms also need to be identified. The understanding of these pathways would open the way to elucidating the mechanisms by which neonicotinoids impact the circadian rhythms of honey bees, and suggest possible means to limit their disruptive impact.

## Materials and Methods

### Honey bees

Honey bees (*Apis mellifera ligustica*) were maintained in standard Langstroth 10-frame boxes (Kelly Beekeeping, Clarkson, KY) on the Vanderbilt campus. All bees used for experiments came from healthy colonies with a mated queen. Vanderbilt does not use neonicotinoid pesticides on its property, and forager honey bees from our hives did not show detectable baseline levels of thiamethoxam or clothianidin, the two neonicotinoids that we used in our study (Supplemental Figure 1).

### Locomotor activity monitoring experiments

Foragers were caught outside the hive entrance upon their return from foraging trips, confirmed by the presence of pollen on their hind legs. Foragers were individually housed in plastic tubes containing either dosed bee candy (ground white cane sugar and honey) containing a specified amount of thiamethoxam, clothianidin, or control bee candy with no pesticide. Tubes were loaded into Locomotor Activity Monitors (LAMs, Trikinetics) and housed in an environmental chamber with a controlled light:dark cycle, constant temperature of 35°C and humidity of 60%. A filter paper wick supplied water from a reservoir to each tube^19^. Data recording (infrared beam interruptions) began immediately after loading the monitors, and data beginning with the first full day of activity was used for analysis. Experiments of 12:12 light:dark cycle (LD) for 4 days, followed by 4 days of constant darkness (DD), took place in the spring/summer of 2017. Experiments of 250 lux constant light (LL) and constant darkness (DD) were performed during the spring/summer of 2019. In the case of the LL experiments, bees were placed into the incubator during the daytime and lights were not turned off for the duration of the experiment. For DD, foragers were placed in the incubator in the daytime and lights were turned off at 19:00 CDT that day and not turned back on for the duration of the experiment. For the LD experiments, locomotor activity rhythms and sleep data were analyzed from bees that survived in the activity monitors a minimum of 4 days, and up to 8 days of data were used. For the DD and LL experiments only bees that survived 5 full days of the experiment were used for data analysis.

### Behavioral metrics (data analysis)

Locomotor activity rhythms are displayed as double plotted actograms (Figure 1A-D). Free-running period calculations were calculated using autocorrelation analysis (Figure 1L, Supplementary Figure 5) generated using the previously published MATLAB toolboxes^45^. Individuals were scored as rhythmic or arrhythmic by a combination of eye inspection of each activity record, the absence of a significant peak in chi-square periodogram, and evaluation of the waveform (showing a significant and cyclical pattern) of the correlogram produced by the autocorrelation analysis (Figure 1E-F, Figure 2). The individual activity onset and offsets for each day were obtained using ClockLab (Actimetrics) with manual corrections as needed (Figure 1 I-L). Average activity duration was calculated by subtracting the onsets from the offsets of each day (offset time – onset time) and then averaging (Supplemental Figure 4A,B). The proportion of activity occurring in the dark was calculated by dividing the number of infrared beam breaks recorded throughout the dark phase of a particular day by the number of beam breaks recorded throughout both the light and dark phases of that day. For each bee, all days were then averaged and these values grouped by treatment (Supplemental Figure 4C,D). Total activity counts for LD portion of the experiment shown as total activity counts over 4 days of light: dark cycle (Supplementary Figure 4 E,F).

Sleep was analyzed from the activity data collected from the LAMs. Several studies have shown that lack of movement (beam interruptions) for five consecutive minutes serves as a reliable measure for sleep in honey bees^29,44,45^. Consistent with these studies we defined sleep as 5 min or longer bouts of inactivity. Data was analyzed using a custom toolbox for MATLAB that was originally developed to measure sleep in flies^31^. For each individual, we calculated sleep in 30 min bins, total sleep duration throughout the day, and the number of average sleep episodes in 24 hrs. (Figure 3, Supplemental Figure 6).

### LC-MS/MS Analysis

Sample analyses were carried out by the Vanderbilt Mass Spectrometry Core Lab using a *Vanquish* ultrahigh performance liquid chromatography (UHPLC) system interfaced to a *Q Exactive HF* quadrupole/orbitrap mass spectrometer (Thermo Fisher Scientific) equipped with an Ion Max API source, a HESI-II electrospray probe, and a 50 um ID stainless steel capillary. Chromatographic separation was performed with a reverse-phase Acquity HSS C18 column (1.7 µm, 2.1×50mm, Waters, Milford, MA) at a flow rate of 300 µl/min. Mobile phases were made up of (A) 0.2 % HCOOH in H_2_O and (B) 0.2% HCOOH in CH_3_CN. Gradient conditions were as follows: 0–1 min, B = 0 %; 1–6 min, B = 0–100 %; 6–6.5 min, B = 100 %; 6.5–7.0 min, B = 100– 0 %; 7.0–10 min, B = 0%. The total chromatographic run time was 10 min; the sample injection volume was 10 µL. A software-controlled divert valve was used to transfer eluent from 0–1.5 min and from 5–10 min of each chromatographic run to waste. The instrument was calibrated weekly over a mass range of *m/z* 195 to *m/z* 1822 with a mixture of caffeine, MRFA, and Ultramark-1621 using the manufacturer’s HF-X tuning software. The mass spectrometer was operated in positive ion mode. The following optimized source parameters were used for the detection of analyte and internal standard: N_2_ sheath gas 40 psi; N_2_ auxiliary gas 10 psi; spray voltage 5.0 kV; capillary temperature 320 °C; vaporizer temperature 100 °C. Quantitation was based on parallel reaction monitoring (PRM) using a precursor isolation width of 2.0 *m/z* and the following MS/MS transitions.

| Analyte | Precursor mass (m/z) | Product mass (m/z) | HCD collision energy | S-Lens RF Level |
| --- | --- | --- | --- | --- |
| Clothianidin | 250.0160 | 169.05450 | 30 | 60 |
| Clothianidin-d<sub>3</sub> | 253.0348 | 172.0733 | 30 | 60 |
| Thiamethoxam | 292.0266 | 211.06510 | 30 | 60 |
| Thiamethoxam-d<sub>3</sub> | 295.0454 | 214.0839 | 30 | 60 |

HCD product ion spectra were acquired in the Orbitrap at a resolving power of 30,000, an AGC target of 2e5, and a maximum injection time of 100 ms. Data acquisition and quantitative spectral analysis were done using Xcalibur version 2.2 sp1 and LCQuan version 2.7.0, respectively. Calibration curves were constructed for clothianidin and thiamethoxam by plotting peak area ratios (analyte / internal standard) against analyte concentrations for a series of seven calibrants, ranging in concentration from 2pg to 1ng. A weighting factor of 1/C_t_^2^ was applied in the linear least-squares regression analysis to maintain homogeneity of variance across the concentration range (% error ≤ 20% for at least four out of every five standards). Foragers were exposed to either 0 ng/g, 25 ng/g or 70 ng/g of thiamethoxam and kept in individual chambers for five days. Individuals from each group were taken at 2hr, 26hrs, 50hrs, 74hrs, and 98hrs after exposure to the pesticide food and flash-frozen in liquid nitrogen and kept at −80°C until processed. Brains were dissected in 100% ETOH chilled with dry ice. Each brain was homogenized in 5% 1:1 MeOH:H_2_O solution, acetonitrile. Internal standards for d_3_-thiamethoxam and d_3_-clothianidin were added at a concentration of 250nM in solution. Homogenate was spun at 3000rpm for 3min after which supernatant was decanted and dried. Dried samples were stored at −20°C until processed for LC-MS. The dried residue was reconstituted in 100 µL of H_2_O/CH_3_OH (3:1), vigorously vortexed, and transferred to 200-µL silanized autosampler vials equipped with Teflon-lined bonded rubber septa.

### Calcium Imaging

Calcium imaging was performed as previously described^46^. Adult male flies (4-7 days post-eclosion) from the cross PDF-Gal4>20XUAS-IVS-GCamp5G 41 were immobilized on 300 mm petri dishes in 3 mL physiological saline containing 128 mM NaCl, 2 mM KCl, 4 mM MgCl2, 35.5 mM sucrose, 5 mM HEPES, and 1.8 mM Ca2+, pH 7.2. Labeled PDF soma were identified under halogen lamp illumination and allowed to rest for 2-3 minutes to photo bleach soma prior to baseline recording. l-LN_v_ neurons we distinguished by soma size and position. Soma were imaged with a 40x water-immersion objective with maximal pinhole aperture on a Zeiss LSM510Meta laser-scanning confocal microscope, using 256 × 256 resolution and a region of interest (ROI) box to achieve ∼186 msec scans. Baseline fluorescence was recorded for 250 frames, followed by direct application of given drug, with a main recording for 1000 total frames. To confirm that explants were viable even when no response to treatment/vehicle was observed, 50μL of 2.5M KCl was applied after 1000 frames to confirm a normal calcium response to high potassium.

## Author Contributions

M.C.T. and D.G.M. conceived of the project. M.A.G.G. and M.C.T. performed pesticide/honey bee behavioral experiments. M.A.G.G. and M.C.T. performed honey bee LL experiments, DD experiments and data analysis. C.A.D, M.C.T., and M.A.G.G., performed *Drosophila* Ca^2+^ imaging experiments. M.C.T, M.A.G.G., C.A.D., D.M., and D.G.M. designed experiments. D.G.M., M.C.T. and M.A.G.G. wrote the paper. J.A.R., K.B., and D.M. provided editing, comments, and experimental guidance.

## Acknowledgements

This study was supported by an NSF Graduate Research Fellowship 0909667 and NIH F31 NS096813-02 to M.C.T., NIH R01 GM117650 and a Vanderbilt Discovery grant to D.G.M. The authors would like to thank Emma Rushton, Chad R. Jackson, Tugrul Giray, Terry Jo Bichell, David Bichell and M. Wade Calcutt for discussions. The authors declare no conflicts of interest.

## Notes

### Competing Interest Statement

The authors have declared no competing interest.

