## Supplemental Figures for "Neonicotinoids Disrupt Circadian Rhythms and Sleep in Honey Bees"

**Author Affiliations**:

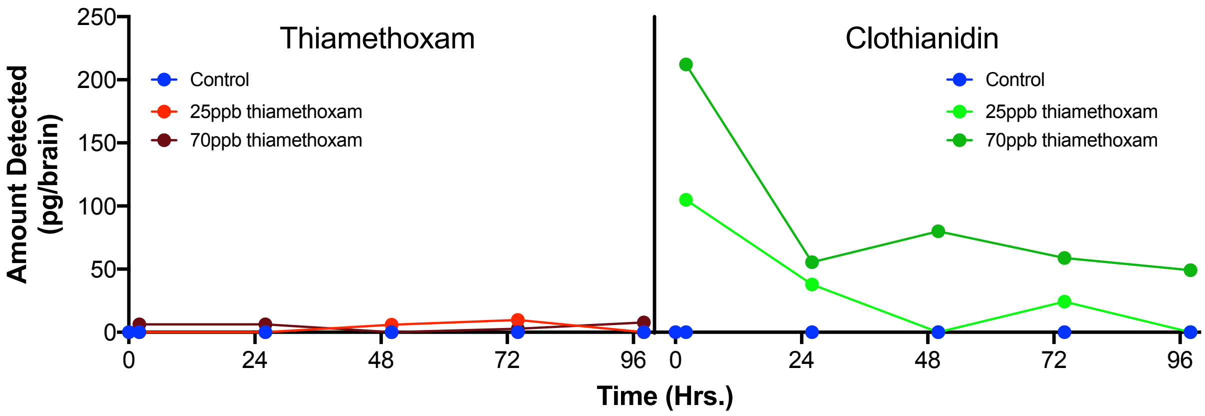


**Supplemental Figure 1:** Time series of LC-MS detection of Thiamethoxam and Clothianidin in the brains of forager honey bees that consumed bee candy dosed with thiamethoxam at 0 ppb (Control), 25 ppb, or 70 ppb.

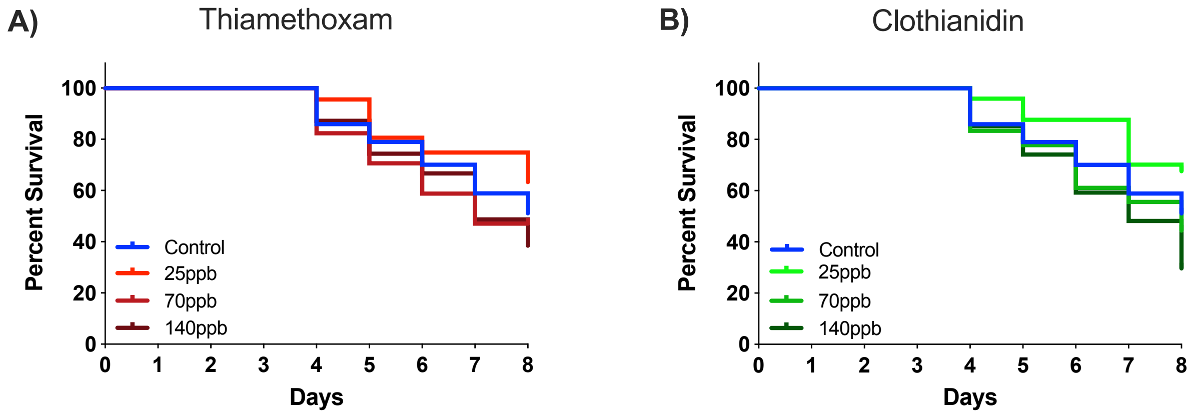


**Supplemental Figure 2**: Survival analysis of bees that contributed data to the LD study. Bee candy dosed with the indicated concentrations of neonicotinoids was available ad libitum for the duration of the experiment. Bees that survived at least 4 days were included in the behavioral analysis. **A)** thiamethoxam exposure at any dose showed no significant differences in mortality from control un-exposed bees on days 4-8. **B)** clothianidin exposure showed no significant increases in mortality compared to unexposed control bees, but did show a significant decrease in mortality at the 30 ppb dose on days 4-8 (Log rank Mantel-Cox test: Thiamethoxam $X^2 = 4.5$ $p = 0.21$; Clothianidin $X^2 = 10.59$ $p = 0.0121$, *).

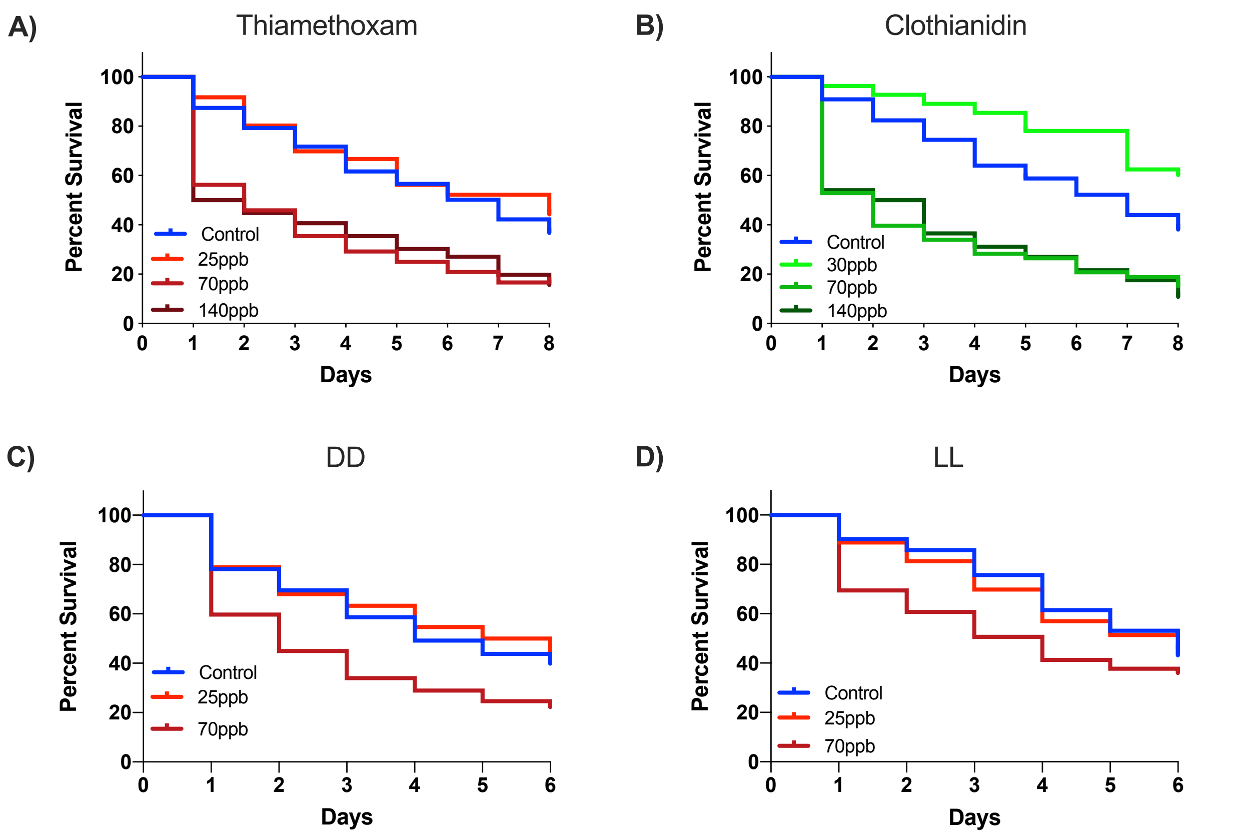


**Supplemental Figure 3**. Survival analysis of all bees that entered behavioral experiments. In general there was an initial significant loss of dosed bees, primarily limited to the first day at the 70 ppb and 140 ppb of thiamethoxam or clothianidin in the food, followed by comparable survivorship to controls on following days. Bees (both control and dosed) not surviving at least 4 days were excluded from the LD experimental analysis, and bees not surviving at least 5 days were excluded from the DD and LL experimental analysis. A) Thiamethoxam or B) Clothianidin in LD experiments (Log rank Mantel-Cox test: Thiamethoxam $X^2 = 44.48$ $p < 0.0001$, ****; Clothianidin $X^2 = 82.39$ $p < 0.0001$, ****). A significant decrease in mortality was observed in the 30 ppb dose of Clothianidin (Log rank Mantel-Cox test: Clothianidin $X^2 = 10.76$ $p = 0.001$, **). Thiamethoxam exposure in both C) DD and LL conditions resulted in significant increases in mortality only for the 70 ppb dose (Log rank Mantel-Cox test: DD 0 vs 70 ppb $X^2 = 17.74$ $p < 0.0001$, ****; LL 0 vs 70 ppb $X^2 = 16.77$ $p < 0.0001$, ****)

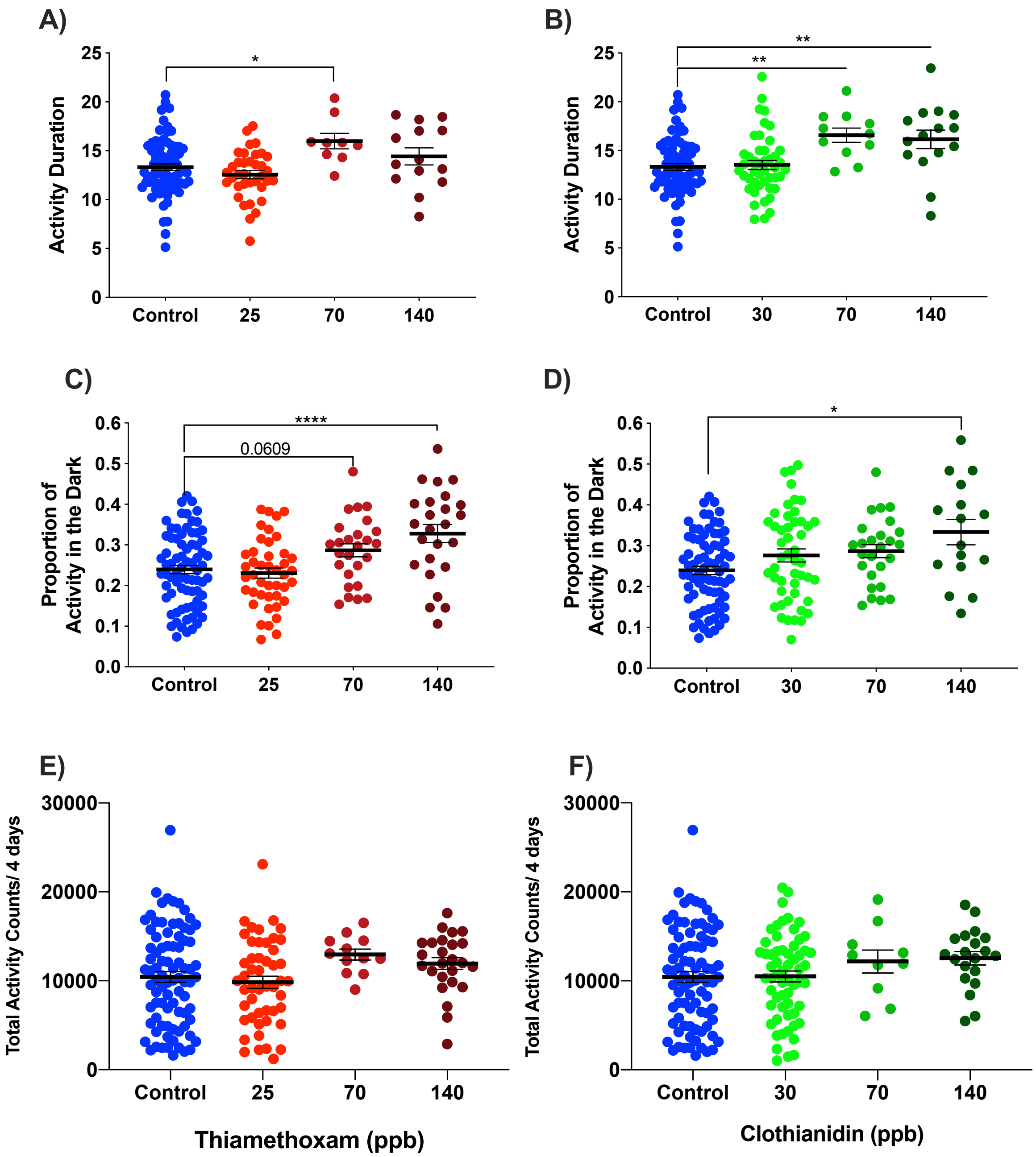


**Supplemental Figure 4**. Chronic exposure to neonicotinoids alters activity duration and the proportion of activity at night but not total activity in foragers. Significant increases in activity duration were observed in both **A)** thiamethoxam (One Way ANOVA, F = 4.257, $p$ = 0.0067, **) and **B)** clothianidin (One Way ANOVA, F = 6.648, $p$ = 0.0003, ***). Dunnett's post-hoc test revealed significant differences between control and pesticide groups as shown. There was also increased activity in the dark phase of the light:dark cycle in both C) thiamethoxam-exposed bees (One Way ANOVA, $F = 8.19$, $p < 0.0001$, ****) and D) clothianidin-exposed bees (One Way ANOVA, $F = 4.817$, $p = 0.0031$, **). Dunnett's post-hoc test revealed significant differences between control and pesticide groups as shown. The observed differences in activity duration and increased activity in the dark were not the result of an overall increase in total activity for either E) thiamethoxam-exposed bees (One Way ANOVA, $F = 2.034$, $p = 0.1111$) or F) clothianidin-exposed bees (One Way ANOVA, $F = 1.313$, $p = 0.2719$).

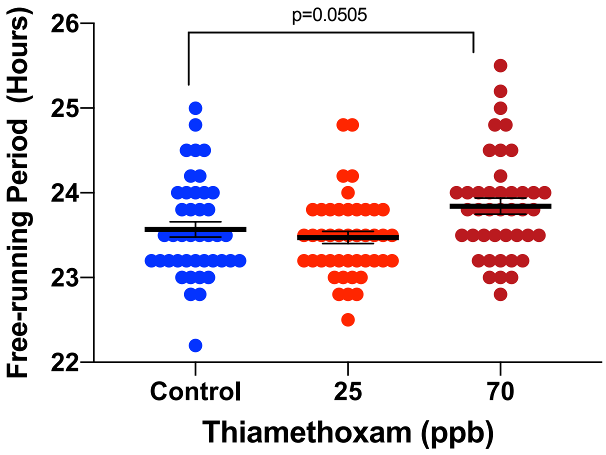


**Supplemental Figure 5.** Altered free-running period after exposure of thiamethoxam in constant darkness. One Way ANOVA revealed significant differences between the groups ($F = 4.979$, $p < 0.0083$, **) as did a test for linear trend (Slope = 0.1369, $p = 0.0274$, *). However, a trend level difference was observed in Dunnett's post hoc test comparing control bees to those exposed to thiamethoxam ($p = 0.0505$).

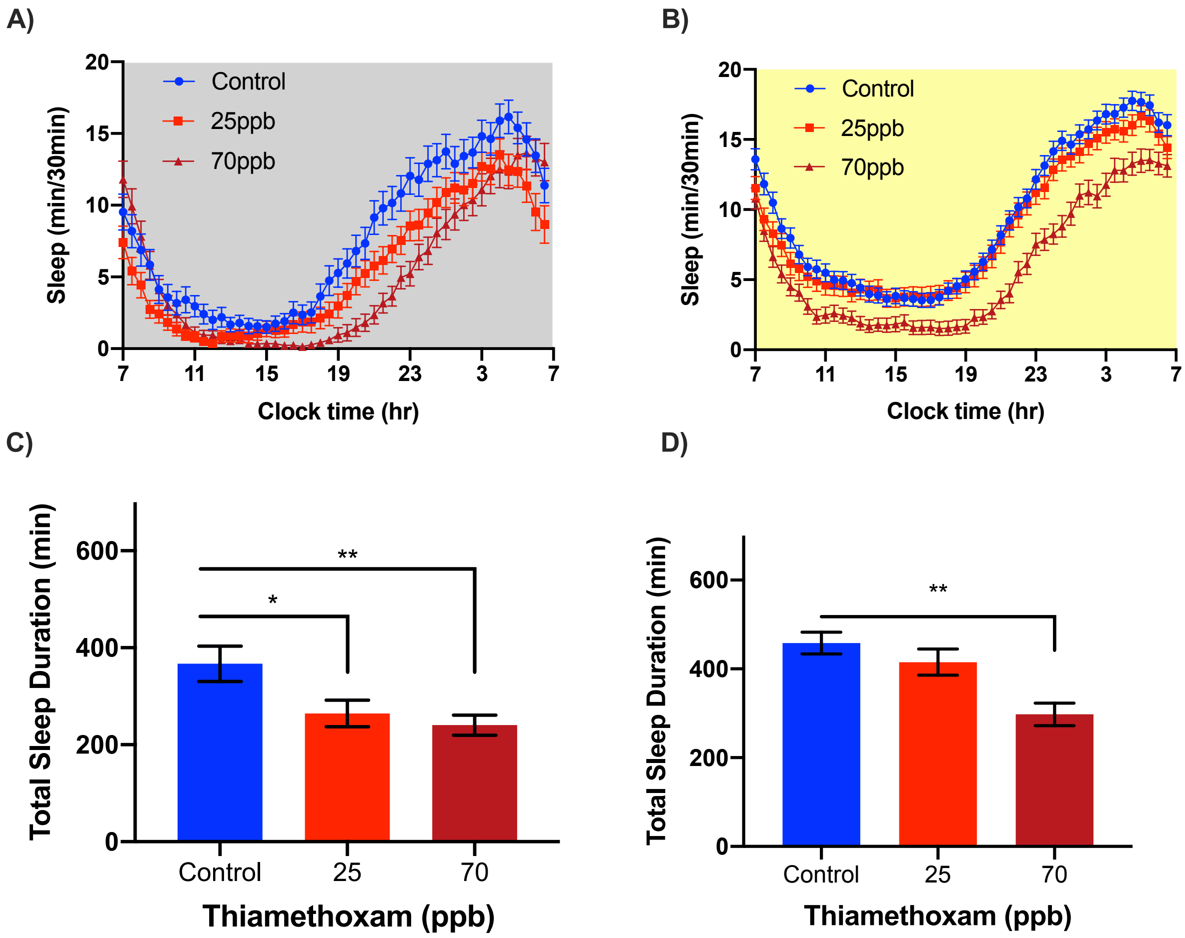


**Supplementary Figure 6.** Thiamethoxam disrupts honey bee sleep under constant conditions**.** Five day average sleep profiles for bees exposed to thiamethoxam at either **A)** DD (Two-way RM ANOVA, Time $p < 0.0001$, ****; Dose $p = 0.0061$, **; Interaction $p < 0.0001$, ****) or **B)** LL (Two-way RM ANOVA, Time $p < 0.0001$, ****; Dose $p < 0.0001$, ****; Interaction $p < 0.0001$, ****). Significant decreases in total sleep duration were also observed in both **C)** DD (One-way ANOVA F = 5.313; $p = 0.0061$, **) and **D)** LL (One-way ANOVA F = 8.644; $p = 0.0002$, ***). Dunnett's post hoc test revealed significant differences comparing controls to each doses as shown.
